# Scaling laws of genome composition and the transition to complex multicellularity

**DOI:** 10.64898/2026.03.02.708964

**Authors:** Rebeca de la Fuente, Wladimiro Díaz-Villanueva, Vicente Arnau, Andrés Moya

## Abstract

Genome architecture reorganizes over evolutionary time to support complex multicellularity without a proportional expansion of coding DNA. We conducted a cross-kingdom comparative analysis using high-quality RefSeq assemblies annotated by the NCBI Genome Annotation Pipeline, restricting the dataset to chromosome-level or complete genomes. Scaling relationships among genome size, gene content, and coding DNA content reveal compositional transitions that distinguish prokaryotic, unicellular eukaryotic, and multicellular lineages. Beyond ∼40 Mb of genic content, coding expansion slows and saturates, indicating compositional constraints that shaped the rise of multicellularity. These results establish scaling laws that quantify how noncoding sequence expansion dominates genome growth in complex eukaryotes.

## 1 Introduction

Biological organization, from molecular networks to whole organisms, obeys certain invariant principles that transcend phylogenetic boundaries. Empirical analyses have shown that diverse biological quantities follow power-law relationships [Banavar et al., 1999]. In physiology, for example, metabolic rate scales with body mass, a pattern observed from bacteria to whales and interpreted as a consequence of energy transport through hierarchical, fractal-like networks [West et al., 1997, Kleiber, 1947]. Allometric scaling relationships are derived in West and Brown [2005] from a general model based on fundamental physical and geometric principles. The West-Brown-Enquist model for allometric scaling, which mechanistically derives the ubiquitous 3/4 power law for metabolic rates, is founded on three physical and geometric constraints common to all biological distribution networks. First, the networks must be space-filling and feature a fractal-like branching geometry to ensure that all cells within the organism’s three-dimensional volume are adequately supplied with resources. Second, the terminal units of the network—such as capillaries, which are the sites of material exchange—are posited to be size-invariant, meaning their dimensions do not change regardless of the organism’s overall body mass, thereby preserving fundamental cellular function. Third, the system has been subject to natural selection for optimization, specifically by minimizing the total energy dissipation (or hydrodynamic resistance) required to transport materials throughout the system. The interplay between these three constraints—fractal geometry, cellular invariance, and energetic optimization—provides a quantitative, first-principles derivation of the 3/4 scaling exponent observed across all domains of life. However, it is worth noting that alternative derivations have been proposed to explain the 3/4 scaling exponent [Kozłowski and Konarzewski, 2004]. For example, in Dodds et al. [2001], the metabolic scaling law was re-examined by minimizing the total impedance of the pulsatile circulatory network instead of the total energy dissipation, leading to an independent derivation of the 3/4 exponent for the metabolic rate. Furthermore, the universal validity of the 3/4 exponent has been questioned by numerous empirical studies demonstrating considerable variation in the data, where scaling exponents oscillate between 2/3 and 1 Glazier [2005]. Therefore, it has been proposed that this observed variation in the exponents can be explained by the Metabolic-Level Boundaries hypothesis, pioneered by Glazier [2005], which posits that the exponent systematically varies between these boundaries—rather than being a universal constant—depending on the organism’s physiological state. Despite ongoing theoretical debate and empirical discrepancies regarding the exact value of the scaling exponent, the broad consensus remains that fundamental scaling laws dominate the organization, function, and evolution of biological systems. These power-law relationships suggest that life, from its most basic molecular networks to the largest ecosystems, operates under stringent universal constraints imposed by physics, geometry, and the optimization of resource transport.

Recent comparative analyses of more than 100 fully sequenced genomes have revealed that the functional composition of genomes follows remarkably regular scaling laws across evolutionary lineages. Using high-level Gene Ontology categories derived from InterPro annotations, [van Nimwegen, 2003] showed that the number of genes in many functional classes scales as a power law with total genome size. These power-law relationships are robust across annotation methods and genome datasets, and their exponents differ systematically between biological processes. For example, metabolic genes scale approximately linearly with genome size, whereas transcription factors scale super-linearly—doubling faster than genome size in bacteria—implying that regulatory complexity expands disproportionately as genomes grow. This disproportionate increase in regulatory complexity is often considered an adaptive outcome. However, it may also emerge as a byproduct of non-adaptive processes, such as genetic drift in small populations [Lynch, 2007]. Conversely, categories related to DNA replication, protein biosynthesis, or the cell cycle scale sub-linearly, indicating tighter evolutionary constraints. A simple evolutionary model based on gene duplication and deletion suggests that these scaling behaviors arise from universal ratios of effective duplication rates across functional classes. These scaling laws suggest that evolution, despite its historical contingency, operates within a constrained phase space shaped by energetic, informational, and physical limits. In this context, the replication of power-law patterns at the molecular level suggests the possibility that “laws of genomic evolution” may exist, comparable in status to those in physics [Koonin, 2011].

While prokaryotic genomes grow mainly by the duplication and addition of new coding sequences, eukaryotic genomes exhibit a pronounced excess of noncoding DNA, regulatory elements, and introns—features intimately tied to the evolution of complex multicellularity [Lynch and Conery, 2003, Elliott and Gregory, 2015]. Recent large-scale analyses have refined and expanded these observations. Molina and van Nimwegen [2009] demonstrated that scaling exponents of functional categories are remarkably conserved across prokaryotic clades, implying universal organizational rules. In contrast, other studies revealed that these exponents can shift across kingdoms, unveiling evolutionary transitions in functional scaling tied to major physiological innovations [Xu et al., 2006, Konstantinidis and Tiedje, 2004]. These results echo the pattern of broken universality seen in physical systems approaching critical points, where continuous variables undergo abrupt reorganizations. In this context, Ferrada [2025] has recently shown that the emergence of eukaryotes can be described as an algorithmic phase transition: a continuous shift in the relationship between gene length and the fraction of noncoding DNA, marking a fundamental change in the way genomes encode and process information. This transition corresponds to a critical threshold ( 1.5 kb) at which intragenic noncoding sequences abruptly appear, coinciding with the onset of complex regulation and multicellular potential. Their model frames genomic evolution as a computational process in which noncoding regions act as algorithmic innovations that enhance the information capacity of genomes.

Building on this theoretical lineage, we analyze genome composition across prokaryotic and eukaryotic lineages to uncover scaling laws linking genome size, coding content, and gene content. Our results reveal a continuous yet nonlinear transition: coding expansion saturates beyond a characteristic threshold ( 40 Mb), delineating a compositional boundary between unicellular and multicellular architectures. We formalize this behavior through a smooth crossover between linear and sublinear regimes, unifying molecular and organismal scaling within a single quantitative framework. Together, these findings suggest that genome evolution is governed by universal constraints analogous to those driving physical criticality—constraints that couple energy, information, and structure across biological scales. The transition to complex multicellularity thus emerges not merely as a historical contingency, but as the inevitable consequence of scaling limits intrinsic to genomic organization.

## 2 Materials and Methods

### 2.1 NCBI RefSeq dataset

Genomic attributes were derived from high-quality, complete genome assemblies retrieved from the NCBI RefSeq repository. Annotation data for each genome assembly was extracted from GFF3 files and structured into tab-delimited tables. Each entry corresponds to a biological feature—such as a gene, mRNA, or CDS—while the associated columns provide metadata including unique IDs, genomic coordinates, and functional roles. The data are structured hierarchically to reflect the parent–child relationships between genes, transcripts (mRNA), and coding sequences (CDSs). In this architecture, each gene is linked to various mRNA transcripts that represent the alternative splicing variants. These transcripts are further associated with specific CDSs, which define the translated regions of the mRNA. Collectively, the CDSs linked to a particular transcript constitute the spliced mRNA, thereby characterizing a unique protein isoform.

Derived from these files, we quantified three genomic variables: genome size (*G*), gene content (*S*), and total coding DNA (*C*). Each metric was derived from the start and end coordinates of the annotated elements. Genome size was defined as the total number of nucleotides in the entire assembly. Gene content was calculated by aggregating all nucleotides located within annotated gene boundaries, while the coding region was determined by summing the nucleotides belonging to at least one CDS.

While our data collection aimed to encompass the entire tree of life, the final sampling was determined by the availability of high-quality whole-genome annotations, yielding a dataset of 694 eukaryotic species. To ensure data quality and consistency, we applied a multi-step filtering process to the available dataset. First, we selected only genomes that the NCBI RefSeq annotation pipeline had processed. Second, the scope was limited to assemblies reaching the “chromosome” or “complete genome” levels to guarantee structural integrity. Third, the dataset was refined to include only major taxonomic groups containing at least 20 species. The set of annotation files was retrieved in November 2024.

### 2.2 Comparative genomic analysis

Phylogenetic relationships for species satisfying the aforementioned criteria were mapped using NCBI Taxonomy tools [Sayers et al., 2023] and subsequently visualized using iTOL [Letunic and Bork, 2021]. We restricted our eukaryotic analysis to taxonomic ranks—specifically kingdom, phylum, and class—represented by at least 20 distinct species. This selection process identified several major clades: 133 mammals, 77 birds, 169 fish, 187 arthropods, 128 plants, 130 fungi, and 53 unicellular eukaryotes. In our dataset, fungi are predominantly unicellular eukaryotes and share a genomic composition similar to that of protists. Regarding the other biological domains, we randomly sampled 400 archaea and 400 bacteria from the vast number of available NCBI annotations. To account for evolutionary relationships, a phylogenetic tree based on common ancestry was generated using the TimeTree platform (timetree.org, [Kumar et al., 2022]) and exported in Newick format. Due to the availability of taxonomic records, the final phylogeny represents a representative subset of our dataset, comprising 170 archaea, 213 bacteria, 35 unicellular eukaryotes, 79 fungi, 110 plants, 116 arthropods, 163 fish, 70 birds, and 122 mammals.

To account for phylogenetic non-independence, relationships between genomic variables were evaluated using Phylogenetic Generalized Least Squares (PGLS) regressions. This approach incorporates shared ancestry into the statistical framework, preventing phylogenetic structure from confounding trait correlations. Model performance and evolutionary constraints were assessed using the scaling coefficient (*β*), the adjusted coefficient of determination 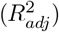, and Pagel’s *λ* to quantify phylogenetic signal. Statistical significance was determined at *p* ≤ 0.05 threshold.

To evaluate relative dispersion across variables with different scales, we calculated the coefficient of variation 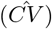, defined as the ratio of the standard deviation (*s*) to the sample mean 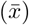, expressed as

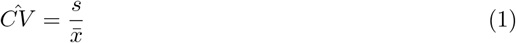

The coefficient of variation 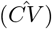 serves as a scale-invariant metric for comparing dispersion across datasets of varying magnitudes. In this context, 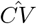values reflect the degree of evolutionary constraint, with higher values indicating genomic heterogeneity and lower values indicating stabilized or constrained traits. Furthermore, comparing these variability regimes across lineages reveals whether genomic heterogeneity follows universal scaling laws or is shaped by taxon-specific evolutionary pressures. To evaluate these inter-variable dynamics, we introduce a relative variability metric defined as the ratio between the coefficients of variation of different features:

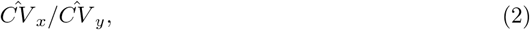

where 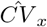 is the coefficient of variation of *x*. These ratios provide a scale-invariant assessment of the relative dispersion among genomic traits, facilitating direct comparison of their variability regimes within and across taxonomic lineages.

Finally, to quantitatively evaluate the scaling relationships, the equations *C* = *aG*^*γ*^, *C* = *aS*^*γ*^ and *S* = *aG*^*γ*^ were fitted independently for each taxonomic group using a GLS regression on log-transformed variables, linearizing the power-law relationship. A global relationship encompassing all species across all taxonomic groups was parameterized using the minimum scaling exponent, *γ*, observed within each taxonomic group. This exponent was incorporated into the general model 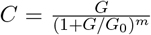, where the parameter *m* is derived as 1 − *γ*. This mathematical model captures well the scaling dynamics observed among various genomic variables. Specifically, the model accurately characterizes the previously described scaling relationship between coding content and genome size, as well as between coding and total gene content. The threshold *G*_0_, obtained by fitting the model to all taxonomic groups, defines the transition from linear to sublinear scaling. This parameter allows the dataset to be characterized into two distinct regimes based on its genomic composition.

## 3 Results

### 3.1 Comparative analysis across taxonomic groups

The prokaryotic genome regime includes bacteria and archaea, whose genome sizes range from 1 Mb to 7Mb and are characterized by compactness, with minimal noncoding DNA. We observe a second regime for the unicellular eukaryotes, with genome sizes spanning approximately one order of magnitude larger than those of prokaryotes, from 1 Mb to 60 Mb. This category includes protists and fungi. Finally, a third regime is observed in multicellular organisms, with genome sizes spanning two orders of magnitude larger than those of unicellular eukaryotes, from 150 Mb to 4500 Mb. Within this regime, we can further distinguish subcategories: plants, fish, and arthropods exhibit genome sizes ranging from 150 Mb to 4500 Mb, spanning a highly diverse range. On the other hand, birds and mammals show a more restricted range. Birds have genome sizes constrained between 1000 Mb and 1500 Mb, and mammals from 2000 Mb to 3400 Mb.

As previously reported in de la Fuente et al. [2025], the percentage of the genome composed of genes exhibits a downward trend from prokaryotes to mammals, with plants characterized by an exceptionally high proportion of intergenic DNA. In bacteria and archaea, gene content accounts for approximately 85% of the genome, dropping to 60% in fungi, unicellular eukaryotes, arthropods, and fish. This proportion is lower in homeothermic vertebrates, ranging from 53% in birds to 45% in mammals. Notably, the gene percentage in plants drops to roughly 25%, sharply diverging from patterns observed in other taxa (see Table 1).

**Table 1.**
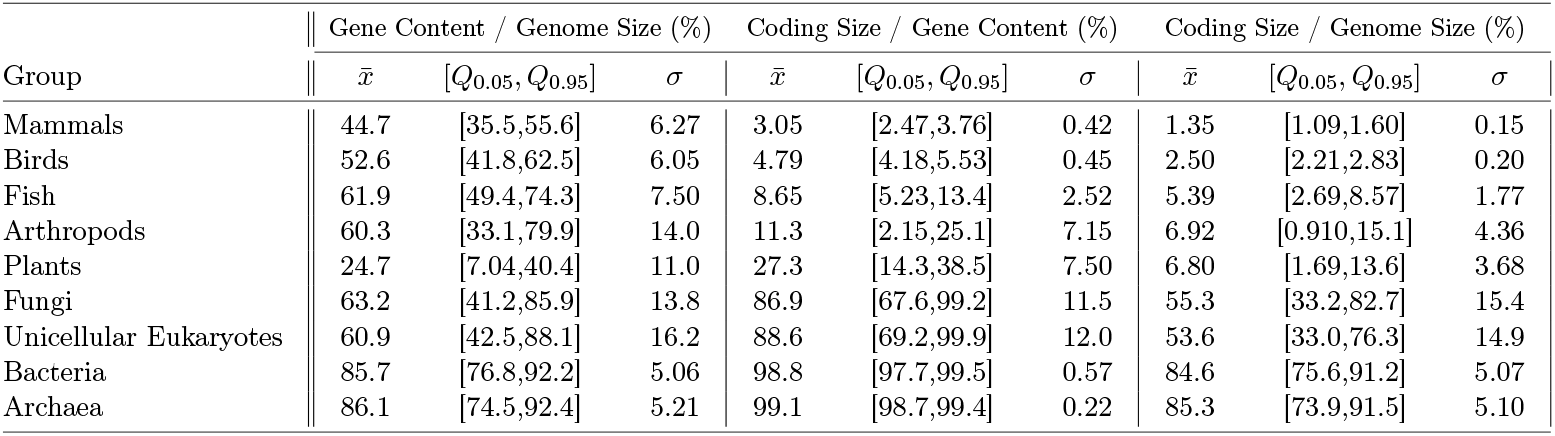
Summary statistics for the percentage of gene content relative to genome size (Gene Content / Genome Size (%)), the percentage of coding relative to gene size (Coding Size / Gene Content (%)), and the percentage of coding relative to genome size (Coding Size / Genome Size (%)). The table includes the mean 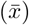, the interpercentile range ([*Q*_0.05_, *Q*_0.95_]), and standard deviation (*σ*) for each group.

| Group | Gene Content / Genome Size (%) |  |  | Coding Size / Gene Content (%) |  |  | Coding Size / Genome Size (%) |  |  |
| --- | --- | --- | --- | --- | --- | --- | --- | --- | --- |
| | $\bar{x}$ | $[Q_{0.05}, Q_{0.95}]$ | $\sigma$ | $\bar{x}$ | $[Q_{0.05}, Q_{0.95}]$ | $\sigma$ | $\bar{x}$ | $[Q_{0.05}, Q_{0.95}]$ | $\sigma$ |
| Mammals | 44.7 | [35.5,55.6] | 6.27 | 3.05 | [2.47,3.76] | 0.42 | 1.35 | [1.09,1.60] | 0.15 |
| Birds | 52.6 | [41.8,62.5] | 6.05 | 4.79 | [4.18,5.53] | 0.45 | 2.50 | [2.21,2.83] | 0.20 |
| Fish | 61.9 | [49.4,74.3] | 7.50 | 8.65 | [5.23,13.4] | 2.52 | 5.39 | [2.69,8.57] | 1.77 |
| Arthropods | 60.3 | [33.1,79.9] | 14.0 | 11.3 | [2.15,25.1] | 7.15 | 6.92 | [0.910,15.1] | 4.36 |
| Plants | 24.7 | [7.04,40.4] | 11.0 | 27.3 | [14.3,38.5] | 7.50 | 6.80 | [1.69,13.6] | 3.68 |
| Fungi | 63.2 | [41.2,85.9] | 13.8 | 86.9 | [67.6,99.2] | 11.5 | 55.3 | [33.2,82.7] | 15.4 |
| Unicellular Eukaryotes | 60.9 | [42.5,88.1] | 16.2 | 88.6 | [69.2,99.9] | 12.0 | 53.6 | [33.0,76.3] | 14.9 |
| Bacteria | 85.7 | [76.8,92.2] | 5.06 | 98.8 | [97.7,99.5] | 0.57 | 84.6 | [75.6,91.2] | 5.07 |
| Archaea | 86.1 | [74.5,92.4] | 5.21 | 99.1 | [98.7,99.4] | 0.22 | 85.3 | [73.9,91.5] | 5.10 |

Furthermore, the proportion of coding sequences within genic regions declines dramatically across the tree of life. While coding DNA constitutes approximately 99% of the gene content in prokaryotes, this value falls to 3% in mammals. In unicellular eukaryotes and fungi, this proportion remains high, accounting for approximately 87% of the gene content. This trend of increasing non-coding genic regions is even more pronounced in multicellular organisms: coding fractions drop to 27% in plants and reach minimal levels in vertebrates, with 9% in fish, 5% in birds, and a mere 3% in mammals. These findings highlight a substantial expansion of non-coding DNA within the genic architecture of complex animals, peaking in the mammalian lineage (see Table 1).

PGLS analyses demonstrate that while genomic correlations remain statistically significant across all studied taxa, the model fits exhibit substantial lineage-specific heterogeneity (see Table S1 in the Supplementary File 1 and Figs. S1-S6 of Supplementary File 2). In prokaryotic lineages (Bacteria and Archaea), the relationships between coding content, gene content, and total genome size are nearly isometric, with the model explaining over 90% of the observed variance 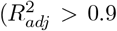 in all cases). These correlations remain high in unicellular eukaryotes and fungi, albeit with slightly lower coefficients than in prokaryotes. In contrast, multicellular lineages exhibit broader dispersion, characterized by lower 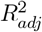 values that suggest a relative decoupling between genome size and functional content. Specifically, the correlation between coding sequences and total genome size is highly predictive in unicellular eukaryotes 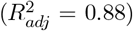, but shows greater dispersion among fungal lineages 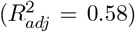. Within multicellular taxa, the model’s predictive power decreases substantially, with 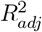 values of 0.41 for plants, 0.28 for fish, 0.24 for mammals, 0.32 for birds, and 0.10 for arthropods. The association between coding sequences and total gene content remains strong in unicellular eukaryotes 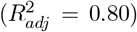 and fungi 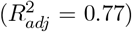. However, this relation is weaker in multicellular organisms, with explanatory power ranging from moderate in birds (0.53) and plants (0.51) to notably lower in fish (0.29), mammals (0.23), and arthropods (0.13). The model’s predictive power for the relationship between gene content and genome size varies substantially across taxa. High 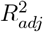 values were observed in fish (0.91), arthropods (0.82), and unicellular eukaryotes (0.76), indicating strong genomic coupling. In contrast, this association weakens in fungi (0.61) and plants (0.41), reaching its lowest levels in birds (0.18) and mammals (0.17).

A notable difference is observed in the relationship between coding proportions with both gene content and genome size. While these variables exhibit high statistical parsimony across several multicellular lineages, they show comparatively lower correlations within prokaryotic groups. The relationship between the coding fraction relative to gene content and the total gene content exhibits a strong negative correlation in higher vertebrates, and a positive correlation in prokaryotes. While 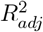 values are relatively low in prokaryotes (Archaea: 0.21, Bacteria: 0.16) and simpler eukaryotes (ranging from 0.23 to 0.32), the model’s predictive power rises sharply in birds 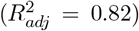 and mammals 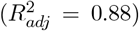, indicating a highly constrained relationship in these lineages. Similarly, the relationship between the coding fraction and total genome size shows the highest goodness-of-fit in mammals 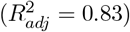 and birds 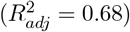, followed by fish 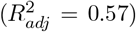 and plants 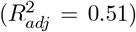. This predictive power diminishes in unicellular eukaryotes (0.44) and fungi (0.29), reaching its lowest value in arthropods (0.13). Notably, this correlation disappears in bacteria 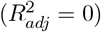, whereas in archaea it remains moderate 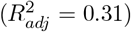. Ultimately, the relationship between relative gene proportion and genome size reveals a consistently negative correlation across nearly all taxa, with the notable exception of Bacteria, where no significant association was detected. The model’s explanatory power was most pronounced in plants 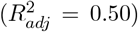, and archaea 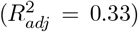, followed by fungi 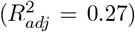 and unicellular eukaryotes 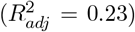. In contrast, the association was considerably weaker in arthropods (0.18), fish (0.13), birds (0.13), and mammals (0.10), suggesting greater genomic variation in these multicellular lineages.

To assess the relative variability of genomic features across taxonomic lineages, we employed a scaling index based on the ratio of their coefficients of variation (see Methods and Supplementary Files 1 and 2). The coefficients of variation for coding and gene content are nearly identical across prokaryotes, fungi, and unicellular eukaryotes, yielding a variability ratio 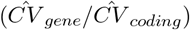 near unity. Conversely, gene content varies more substantially than coding content within multicellular taxa, with arthropods displaying the lowest relative variability 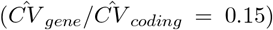. Our results reveal a fundamental shift in genomic architecture between domains. In prokaryotes, the coding percentage exhibits minimal variation despite significant fluctuations in total gene content, resulting in a variability ratio 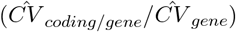 near zero. In contrast, in multicellular organisms, we observe a marked increase in the variability of the coding proportion alongside a reduction in the variance of gene content. Consequently, these two metrics display more comparable degrees of dispersion, driving the ratio toward unity in more complex taxa. Specifically, the ratio increases from a near-zero value in prokaryotes (0.01) to 0.17 and 0.26 in unicellular eukaryotes and fungi, respectively. Thus, the upward trend continues through plants (0.47) and arthropods (0.49), reaching its highest levels in vertebrates, with values of 0.61 for fish, 0.77 for birds, and 0.85 for mammals. A similar pattern emerges for the 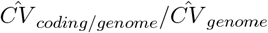. Values increase from the lower levels found in bacteria (0.11) and archaea (0.16) through unicellular eukaryotes (0.27), fungi (0.37), and arthropods (0.34). The ratio reaches its peak in plants (0.49) and especially in vertebrates, with fish, birds and mammals showing values of 0.58, 0.80, and 0.83, respectively. The results also reveal that the variability of gene content is nearly equal to genome size in several lineages, with the ratio of their coefficients of variation 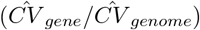 approaching unity in mammals, birds, bacteria, and archaea. However, plants exhibit a distinct pattern in which genome size variability significantly outpaces gene content variability, yielding a minimum ratio of 0.52. Finally, the variability ratio between the gene/genome proportion and total genome size 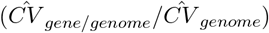 shows a marked increase across taxa. While values are relatively low in prokaryotes (0.11-0.16), arthropods (0.12), and fish (0.21), they rise through unicellular eukaryotes (0.26), fungi (0.29), and plants (0.40). Notably, this ratio exceeds unity in endothermic vertebrates, reaching 1.11 in birds and 1.04 in mammals, indicating that, in these groups, the gene/genome proportion is more variable than genome size itself.

### 3.2 Global scaling patterns across the tree of life

#### 3.2.1 The mathematical model

We empirically derived a mathematical model that fits the relationship between the different genomic variables. As observed in Fig. **??**, prokaryotic genome expansion is primarily driven by an increase in coding sequences. However, as genome size increases, we observe a transition at the origin of complex multicellularity and genome expansion in larger organisms is increasingly dominated by non-coding elements. The evolutionary transition from minimal prokaryotic genomes to huge genomes in the multicellular regime occurs throughout the unicellular eukaryotic regime, which represents an intermediate state. We introduce a mathematical model that describes the genome composition across the distinct evolutionary regimes and captures the transition between them:

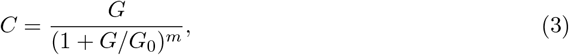

where *C* denotes the coding DNA content and *G* the genome size. This model captures a continuous transition in genome composition, in which where coding content initially scales linearly with genome size but gradually shifts to a sublinear regime as genome expansion incorporates increasing amounts of non-coding DNA. The transition between these two scaling regimes occurs at the evolutionary shift from unicellular to multicellular organisms, leading to saturation effects in the latter regime. The model is characterized by two key parameters: *G*_0_. This critial genomic threshold defines the transition point between regimes, and the scaling exponent *m*, which determines the degree of deviation from linearity in large genomes. When genomes are much smaller than the critical threshold, *G* ≪ *G*_0_, the term *G/G*_0_ becomes negligible. In this case, the equation reduces to *C* ≈ *G*, indicating a linear relationship between coding content and genome size. This regime is characteristic of prokaryotic genomes, where nearly all genomic expansion corresponds to an increase in coding DNA. When genomes are much larger than the threshold, *G* ≫ *G*_0_, the term 1 + *G/G*_0_ dominates *G/G*_0_, so the denominator behaves as (1 + *G/G*_0_)^*m*^ ≈ (*G/G*_0_)^*m*^. Substituting this term into the equation, we obtain the following approximation: 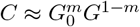. This indicates sublinear scaling, where coding content grows more slowly as genome size increases. In this regime, coding content growth slows down as genome expansion becomes dominated by non-coding sequences. Thus, it exhibits a saturation controlled by the parameter *m*, which governs the transition between linear and sublinear scaling. The parameter can take values in the range of 0 ≤ *m* ≤ 1, modulating the degree of saturation. When *m* = 0, the model recovers a purely linear scaling, meaning that the same scaling law applies across both small and large genomes, and no transition occurs. In this case, genome expansion remains driven primarily by the accumulation of coding sequences, as observed in prokaryotic genomes. For *m* = 1, the transition leads to complete saturation of the coding content, reaching an upper bound. However, an intermediate scenario arises when 0 *< m <* 1, *dC* in which the system undergoes a gradual transition, with the rate of coding expansion progressively slowing as genome size increases. Notably, as genome size increases, the accumulation of noncoding DNA becomes increasingly dominant, leading to asymptotic saturation of coding content. Mathematically, taking the derivative of the equation in the regime of large genomes, we obtain:

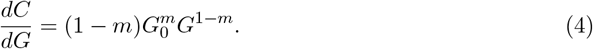

When *m >* 0, the following asymptotic relation holds:

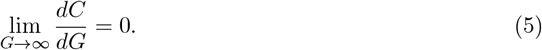

If genome size increases indefinitely, the rate of growth in coding content approaches zero. Thus, the model predicts an asymptotic saturation of coding content as genome size increases.

From the fitted scaling relation, we can derive the conditional probability that a newly added genomic fragment is coding:

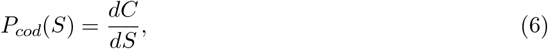

which gives:

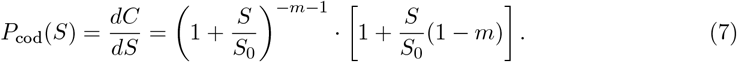

This function defines a size-dependent Bernoulli process governing the genic expansion, where each added fragment is coding with probability *P*_*cod*_(*S*). The overall coding content *C*(*S*) then corresponds to the expected cumulative number of successes of this process. This function decreases with increasing S: as total genic content grows, the probability that newly added fragments are coding decreases. Furthermore, the decay follows a power-like behavior. Thus, it allows *P*_*cod*_(*S*) to be used as a probabilistic decision rule in Monte Carlo simulations to determine whether an added fragment is coding or non-coding.

#### 3.2.2 Scaling patterns

The model describes two genomic regimes, each governed by distinct scaling laws:

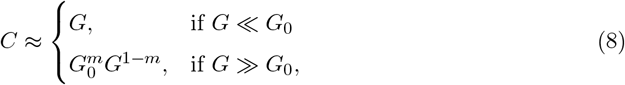

First, the scaling exponent *C* ∼ *G*^*γ*^ was derived for each taxonomic group based on the relation *γ* ≈ 1 − *m* (see Table 4 of Supplementary File 1). All pairwise relationships were found to be statistically significant. Our results demonstrate that coding content scales linearly with genome size in both archaea and bacteria, with scaling exponents close to unity 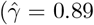 and 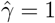, respectively). However, in the unicellular eukaryotic regime, we observe a deviation from this linear scaling, with coding content beginning to grow at a sublinear rate as genome size increases. The estimated exponents are 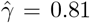 for unicellular eukaryotes and 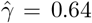 for fungi, reflecting a progressive increase in non-coding sequences. This trend becomes even more pronounced in multicellular organisms, where genome expansion is increasingly driven by the accumulation of noncoding DNA, leading to a sharp decline in the scaling exponent. Specifically, we estimate 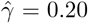 for plants, 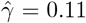 for arthropods, 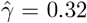 for fish, 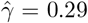 for birds, and 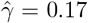 for mammals. These results suggest that the rate of coding expansion is slowing down in multicellular taxa. Second, we estimated the global relationship across all taxa. In this context, the parameter values for the global trend (Eq. 3) were set at 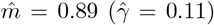 reflecting the scaling exponent of arthropods, which represents the lower bound observed among multicellular groups. Estimating *G*_0_ across the whole dataset yielded Ĝ_0_ = 20 Mb, placing it within the expected unicellular eukaryotic regime (10*Mb <* Ĝ_0_ *<* 100*Mb*). The theoretical model (Fig. 1) aligns robustly with the data, confirming its ability to describe the observed scaling trend.

**Figure 1.**
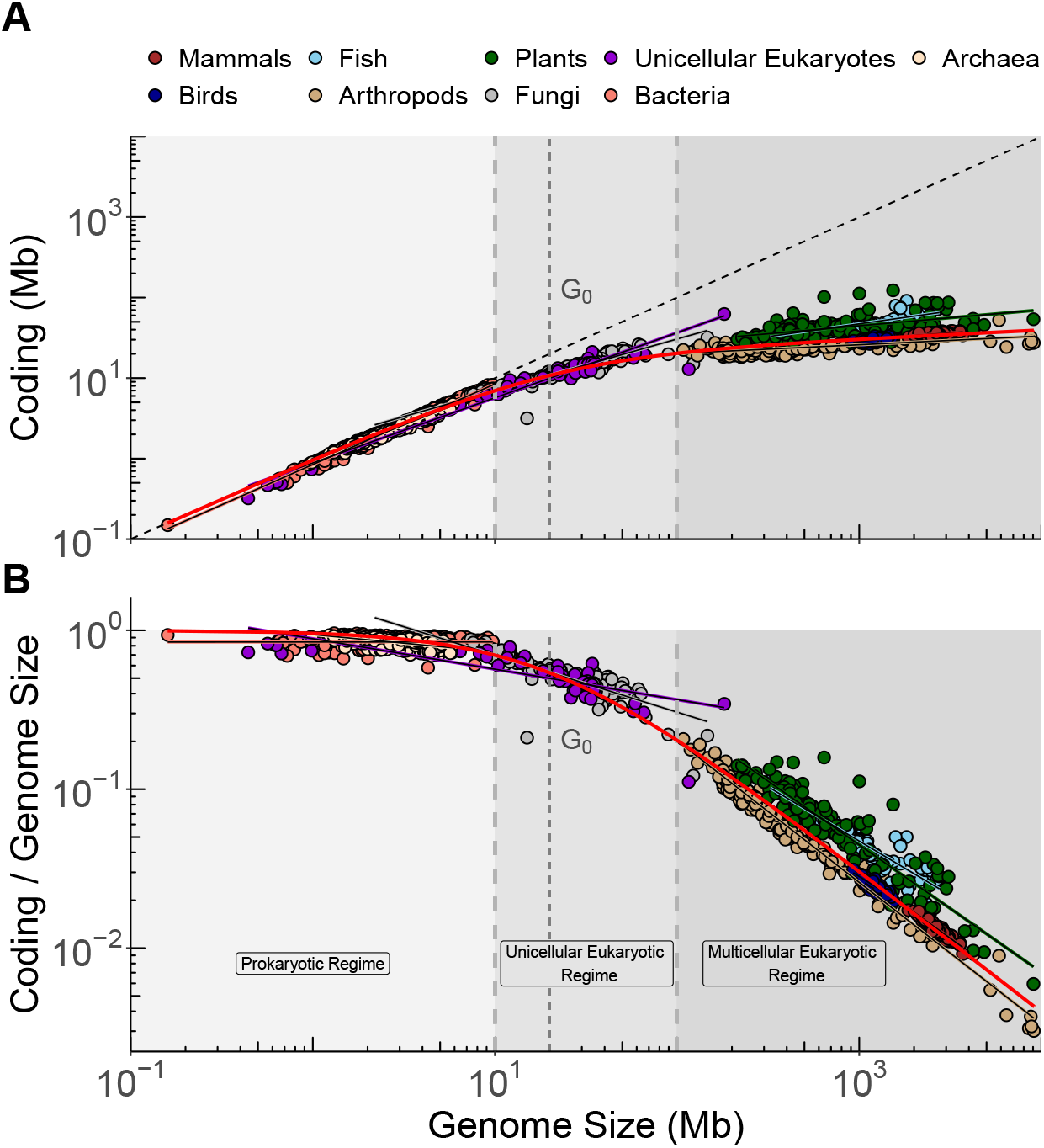
Relationship of genome size with (A) coding DNA content and (B) the proportion of coding DNA. Colored points represent distinct taxonomic groups across prokaryotic (*G <* 10 Mb), unicellular eukaryotic (10 Mb *< G <* 100 Mb), and multicellular eukaryotic (*G >* 100 Mb) regimes. The equation *C* = *aG*^*γ*^ was fitted independently for each taxonomic group. The solid lines represent the best-fit scaling models for each group. The red curve represents the best-fit scaling-law model from Eq. 3. The scaling exponent *m* is approximated as *m* ≈ 1 − *γ*. Here, the scaling exponent is 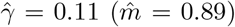, corresponding to the lowest estimated exponent found in arthropods. The transition point was calculated by fitting the model to the complete dataset, yielding Ĝ_0_ = 20 Mb. Note the logarithmic scale on both axes.

The scaling model 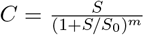, illustrated in Fig. 2(A-B), also captures the relationship between coding and gene content. Consistent with the previous case, the red curve represents the best-fit scaling-law model, which describes a continuous transition from coding-dense prokaryotic genomes to asymptotic coding saturation as genic regions expand in multicellular organisms. First, the best-fit scaling exponent for each group, *C* = *aS*^*γ*^, was estimated (see Table 4 of Supplementary File 1). In prokaryotes, coding content scales almost linearly with the gene content, with a scaling exponent close to 1. In contrast, in multicellular taxa, a progressive saturation of coding is observed. Specifically, we found a scaling exponent of 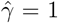 for archaea and bacteria, 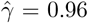 for unicellular eukaryotes, 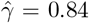 for fungi, 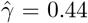 for plants, 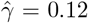 for arthropods, 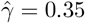 for fish, 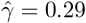 for birds, and 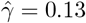 for mammals. We set the scaling exponent to 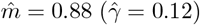 in the global model, corresponding to the lowest estimated exponent reported in arthropods. The transition point was found to be at *S*_0_ = 46 Mb, which falls within the unicellular eukaryotic regime, between *S*^∗^ = 10 Mb and *S*^∗^ = 60 Mb. As observed, the accumulation of coding DNA slows down as the gene content increases.

**Figure 2.**
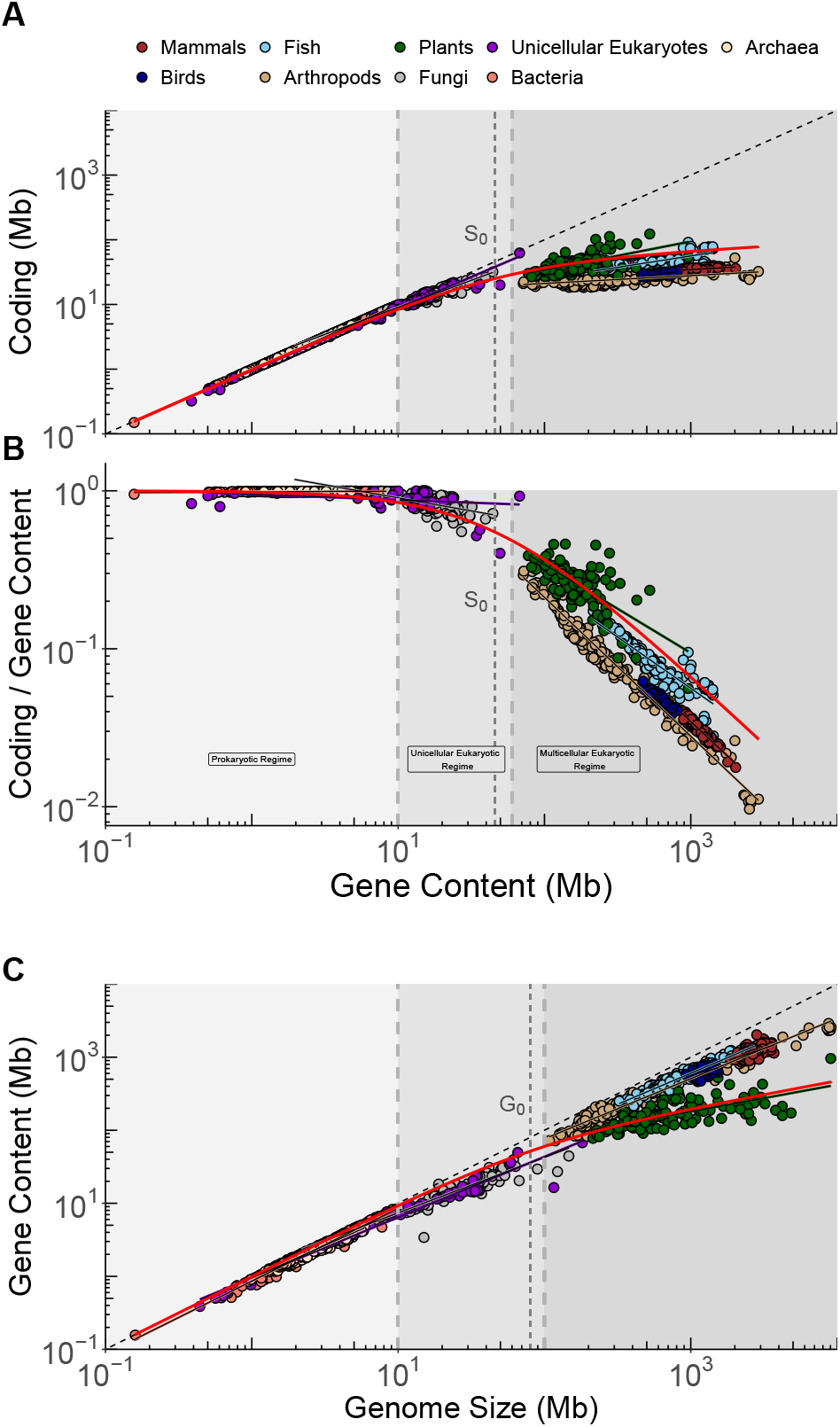
Relationship of gene content with (A) coding DNA content and (B) the proportion of coding DNA in genes. Colored points represent distinct taxonomic groups across prokaryotic (*S <* 10 Mb), unicellular eukaryotic (10 Mb *< S <* 60 Mb), and multicellular eukaryotic (*S >* 60 Mb) regimes. The equation *C* = *aS*^*γ*^ was fitted independently for each taxonomic group. The solid lines represent the best-fit scaling models for each group. The red curve represents the best-fit scaling-law mode. Here, the scaling exponent is 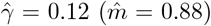, which corresponds to the lowest estimated exponent, found in arthropods, and the transition point was estimated by fitting the model to the complete dataset, which was found at Ĝ_0_ = 46 Mb. (C) Relationship of the gene content with genome size across prokaryotic (*G <* 10 Mb), unicellular eukaryotic (10 Mb *< G <* 100 Mb), and multicellular eukaryotic (*G >* 100 Mb) regimes. The equation *S* = *aG*^*γ*^ was fitted independently for each taxonomic group. The scaling exponents are close to 1 in all taxonomic groups, except for plants, which show an asymptotic saturation pattern. The red curve represents the best-fit given by 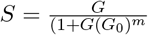, which describes a continous transition from prokaryotic genomes to an asymptotic saturation of the gene content in plants. Here, the scaling exponent is approximated by the scaling-law observed in plants, with 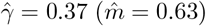. The transition point was estimated from data across prokaryotes, unicellular eukaryotes and plants, yielding *G*_0_ = 80 Mb. Note the logarithmic scale on both axes.

In contrast, the expansion of gene content with increasing genome size scales almost linearly across all taxa, except plants. The scaling exponent of this trend was also estimated for each taxonomic group. As shown in Fig. 2(C), we find that archaea 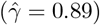 and bacteria 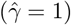 exhibit near-linear scaling, indicating that genome expansion in these groups is proportional mainly to gene acquisition. Similarly, unicellular eukaryotes 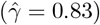 and fungi 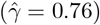 exhibit approximately linear trends, with only slightly deviations from unity. Among multicellular groups, we observe that arthropods 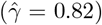 and fish 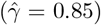 also follow an almost linear scaling, suggesting that genome growth in these taxa remains tightly linked to increases in gene content. In contrast, mammals 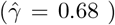 and birds 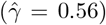 display a more pronounced deviation. However, plants are the only group exhibiting a saturation-like behavior, with a significantly lower scaling exponent 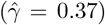, reflecting key differences in genome evolution strategies between plants and animals. In the general model, the scaling exponent was fixed at *m* = 0.63, consistent with the scaling observed in plants. The transition point, *G*_0_, was estimated from data points on prokaryotes, unicellular eukaryotes, and plants and found to be at Ĝ_0_ = 80 Mb. In this case, the model also exhibits a strong fit to the data, suggesting that gene content in plants follows a power-law decay as genome size increases. Furthermore, this result indicates that gene accumulation slows down progressively in plant genomes, although at a moderate rate.

### 3.3 Discussion

The genomic scaling laws identified in this study provide a quantitative framework for understanding the transition from prokaryotic simplicity to complex multicellularity. Our results demonstrate that genome evolution is not merely a stochastic process but is governed by universal structural constraints. As genome size expands, the relationship between coding content and total genomic material undergoes a fundamental shift from linear, coding-dense regime to an asymptotic sublinear saturation. A central finding of our analysis is the empirical derivation of the critical genomic threshold, *G*_0_ = 20 Mb. This value accurately delineates the boundary between unicellular and multicellular architectures. Below this threshold, characteristic of prokaryotes, genome expansion is nearly isometric (*γ* = 1), meaning growth is driven almost exclusively by the addition of new coding sequences. However, as genome size increases, they enter a regime in which coding expansion slows significantly, and noncoding DNA accumulation becomes the dominant driver of genome growth.

The lineage-specific heterogeneity observed in our PGLS analyses suggest that, while universal laws exist, evolutionary pressures vary across the tree of life. Multicellular taxa exhibit a marked decoupling between genome size and functional content, in contrast to the high predictive power observed in prokaryotes. Among multicellular taxa, arthropods exhibit the lowest scaling exponent (*γ* = 0.11), which we utilized to parametrize the global saturation limit of the model (*m* = 0.89). This highlights a severe constraint on coding expansion in complex animal lineages.

A key strength of the proposed model is its ability to characterize not only the relationship between coding DNA and total genome size but also the interplay between coding and total gene content. Analogously to the genome-wide trend, the scaling between C and S transitions from a coding-dense regime in prokaryotes to asymptotic saturation as genic regions expand in multicellular organisms. This relationship is particularly revealing when considering the probabilistic nature of genomic growth. As described by the size-dependent Bernoulli process *P*_*cod*_(*S*), the probability that a newly added genomic fragment within a gene is coding decreases as the total genic content increases. In prokaryotes, where *γ* = 1, almost every addition to the gene content results in a corresponding rise in coding DNA. However, in multicellular taxa such as mammals (*γ* = 0.13) and arthropods (*γ* = 0.12), gene expansion is almost entirely driven by noncoding insertions (e.g., introns), leading to the observed sublinear saturation. The fact that the same mathematical framework—with a transition threshold *S*_0_ = 46 Mb situated within the unicellular eukaryotic regime—describes both C(G) and C(S) suggests a common evolutionary constraint. This implies that the “filling” of genomes with non-coding DNA is not a random process, but follows a structured scaling law that governs how information is packed within biological sequences.

Also, unlike most multicellular groups, in which gene content remains tightly linked to genome growth, plants exhibit a unique saturation-like behavior (*γ* = 0.37). This suggests that plants have adopted a distinct evolutionary strategy in which gene accumulation slows as genome size increases, possibly due to unique mechanisms of duplication and intergenic expansion. Analogously to allometric laws in physiology, genomic scaling laws suggest that energetic and informational limits bound the rise of complexity. The transition to complex multicellularity thus emerges as an inevitable consequence of these scaling limits. Future research should investigate whether these mathematical thresholds correspond to specific metabolic costs associated with maintaining and replicating large, non-coding-heavy genomes.

The proposed model allows for the definition of a genomic efficiency ratio, the relationship between adequate coding capacity (C) and total potential capacity (S and G). Our results demonstrate that this ratio follows a universal decay law: while prokaryotes operate near maximum efficiency, the transition to multicellularity shifts potential capacity increasingly toward noncoding regulatory elements. This suggests that the rise of biological complexity is paradoxically linked to a reduction in relative coding density, governed by the scaling parameters *G*_0_ and *m*.

## Data availability

Data has been downloaded from the NCBI database [Kitts et al., 2015]. The list of species under study, code, and the datasets generated in this study are available in Zenodo, at https://doi.org/10.5281/zenodo.

## Acknowledgements

We thank collaborators and institutions that support this research. FISABIO and UAM supported R.d.l.F.M..

